# Parental childhood maltreatment associates with offspring left amygdala volume at early infancy

**DOI:** 10.1101/2023.02.23.529799

**Authors:** Jetro J. Tuulari, Elmo P. Pulli, Eeva-Leena Kataja, Laura Perasto, John D. Lewis, Linnea Karlsson, Hasse Karlsson

## Abstract

**Background:** Childhood maltreatment exposure (CME) and related trauma could be considered some of psychiatry’s greatest public health challenges. CME and early adversity have been associated with increased amygdala volume in exposed individuals. Emerging evidence implies that CME could also affect prenatal development of the offspring.

**Methods:** As part of the FinnBrain Birth Cohort Study, we measured bilateral amygdala volumes from MR images in 76 healthy infants at 2–5 weeks of gestation corrected age and obtained the Trauma and Distress Scale (TADS) questionnaire from both parents. The associations between neonatal amygdala volumes and TADS scores were examined in stepwise regression models.

**Results:** We found that maternal CME associated positively with infant left amygdala volume (p = .045) while the positive association for the paternal trauma score was only marginally significant (p = .099). Similar associations were not observed for the right amygdala. In the exploratory analyses, we used age ranges (0–6, 7–12, and 13–18 years) as estimate of the timing of the CME and included all three time points from both parents using left amygdala volume into the stepwise regression models. We found that maternal TADS scores from 13–18 years of age associated positively with infant left amygdala volumes (p = .008). Correspondingly, paternal TADS scores from 0–6 years of age associated positively with the infant left amygdala volumes (p = .014).

**Conclusions:** Our link the infant offspring amygdala volume with parental CME with some agreement with prior findings, and they also imply links paternal CME to infant amygdala volumes. Amygdala is one of the key brain structures associated with both early life exposures and later psychiatric health, which makes it crucially important to elucidate both the underlying mechanisms and the later relevance of these associations in future studies.

## Introduction

Adverse childhood experiences (ACEs) have long lasting consequences to later health (Hughes et al. 2017). ACEs can be categorized broadly to child maltreatment and life events that may be traumatic. Childhood maltreatment exposure (CME) is highly prevalent. Globally, 12% of adults report childhood sexual abuse, 23% physical abuse, and 36% emotional abuse (Stoltenborgh et al. 2015). CME has well established adverse effect on later mental health but some of its effects may also be transmitted to offspring through maternal physiology during pregnancy (Moog et al. 2018). Intriguing findings point out that CME could be a contributor to fetal brain development (Demers et al. 2022; Khoury et al. 2021; Moog et al. 2018) by programming the maternal physiology (van den Heuvel 2021; Moog et al. 2022). Recent evidence also suggests that paternal CME influences prenatal brain development (Karlsson et al. 2020), potentially via epigenetic changes in the paternal germ line (Ghai and Kader 2021).

Prior research has put much focus on maternal prenatal distress and commonly used questionnaire measures of maternal depressive, anxiety, and stress symptoms (Lautarescu, Craig, and Glover 2020). The amygdala is one of the key structures that has been implicated in animal work related to maternal prenatal distress, and likely has relevance to later child development and health. The amygdalae reach anatomical “maturity” early in development (Bachevalier et al. 1986; Berger et al. 1990; Gabard-Durnam et al. 2018; Humphrey 1968; Kordower, Piecinski, and Rakic 1992; Machado and Bachevalier 2003). Individual differences in early amygdala volume have been linked to maternal pre- and postnatal psychological distress (Supplementary Table 1), although the results have been inconsistent, ranging from positive (Donnici et al. 2021; Groenewold et al. 2022; Wen et al. 2017) to negative (Acosta, Tuulari, et al. 2020; Lehtola et al. 2020; Wang et al. 2018; Wu et al. 2021) with some studies reporting no associations (Hidalgo et al. 2022; Lautarescu et al. 2021; El Marroun et al. 2016; Moog et al. 2021) between maternal pre- or postnatal distress and offspring amygdala volume.

Although the above-mentioned studies in humans do find some parallels to animal studies, the lack of replications and systematic patterns in human studies is evident. This may be due to differences in measurement of both exposures and outcomes, the timing of the exposure. CMEs can underlie adult-age psychiatric symptoms and measures of distress symptoms only might give an incomplete picture. Thus maternal and paternal ACEs, including CME, could be factors that explain the high variability among the studies. To the best of our knowledge, only three previous studies have investigated the associations between maternal CMEs and infant amygdala volumes. Demers et al. (2022) reported a negative association between maternal CMEs and infant bilateral amygdala volumes, and Khoury et al. (2021) showed that the effect between maternal CMEs and right amygdala volume became stronger with increasing age (from 4 to 24 months of age). Moog et al. (2018), on the contrary, did not find any associations between maternal CMEs and infant amygdalae when controlling for pre- and postnatal maternal distress. To our knowledge, no studies have been conducted on the associations of paternal CMEs on infant amygdala volumes.

In the current study, we tested the associations between parental (maternal and paternal) CME with offspring amygdala volumes. Parental CME was measured by the Trauma and Distress Scale (TADS) questionnaire and offspring amygdala volumes from structural magnetic resonance imaging (MRI) soon after birth (minimizing the effect of postnatal exposures). We expected to find negative associations between parental CME scores and amygdala volumes. We did not place a hypothesis on the lateralization of the association.

## Methods

This study was conducted in accordance with the Declaration of Helsinki, and it was approved by the Joint Ethics Committee of the University of Turku and the Hospital District of Southwest Finland (ETMK:31/180/2011). We followed the Strengthening the Reporting of Observational Studies in Epidemiology (<u>STROBE</u>) reporting guideline.

### 2.1. Participants

The participants involved in this study were drawn from the broader FinnBrain Birth Cohort Study (finnbrain.fi; Karlsson et al. 2018). Pregnant women (n□= □3808), their spouses (n□= □2623) and babies to-be born (n□= □3837; including 29 twin pairs) were recruited at gestational week (GW) 12 in Southwest Finland between December 2011 and April 2015. From this broader participant pool, 189 infant-mother dyads were recruited into this study. They were recruited based on willingness to participate and availability of the infant to have an MRI after birth. After explaining the study’s purpose and protocol, written informed consent was obtained from the parent(s). Of these 189 infant participants, 9 did not have any imaging data and 55 had unsuccessful scans or motion artifacts in the MR images, leaving 125 images for further segmentation analyses. Furthermore, 51 were missing parental TADS assessment, leading to a final sample size of 74 triads.

The demographics of the participants are presented in Table 1. Ethnicity was not inquired but the study population was predominantly Scandinavian / Caucasian. Infants were generally scanned two to five weeks after birth, although seven scans were performed after this age, and two before. All infants exceeded 2500□g of weight at birth, and all were term-born. Infant background information was gathered from the Finnish Medical Birth Register (FMBR) kept by the Finnish Institute for Health and Welfare (www.thl.fi). Parental background information was gathered by questionnaires and included monthly income, educational level, and the use of selective serotonin reuptake inhibitors (SSRI), serotonin– norepinephrine reuptake inhibitors (SNRI), alcohol, and tobacco during pregnancy. Obstetric data and maternal pre-pregnancy body mass index (BMI) were retrieved from the FMBR.

**Table 1.**
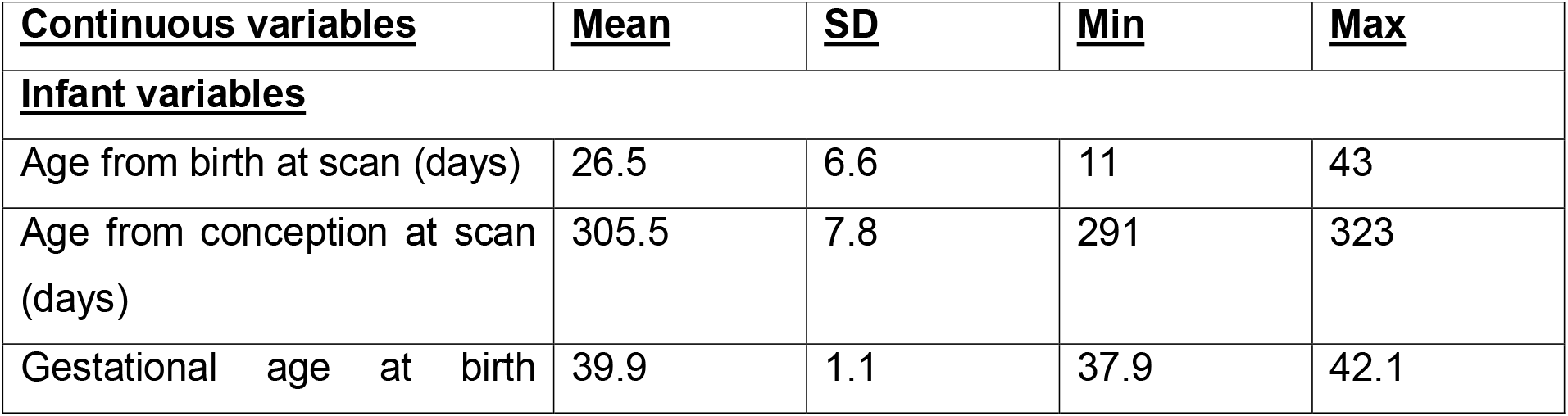
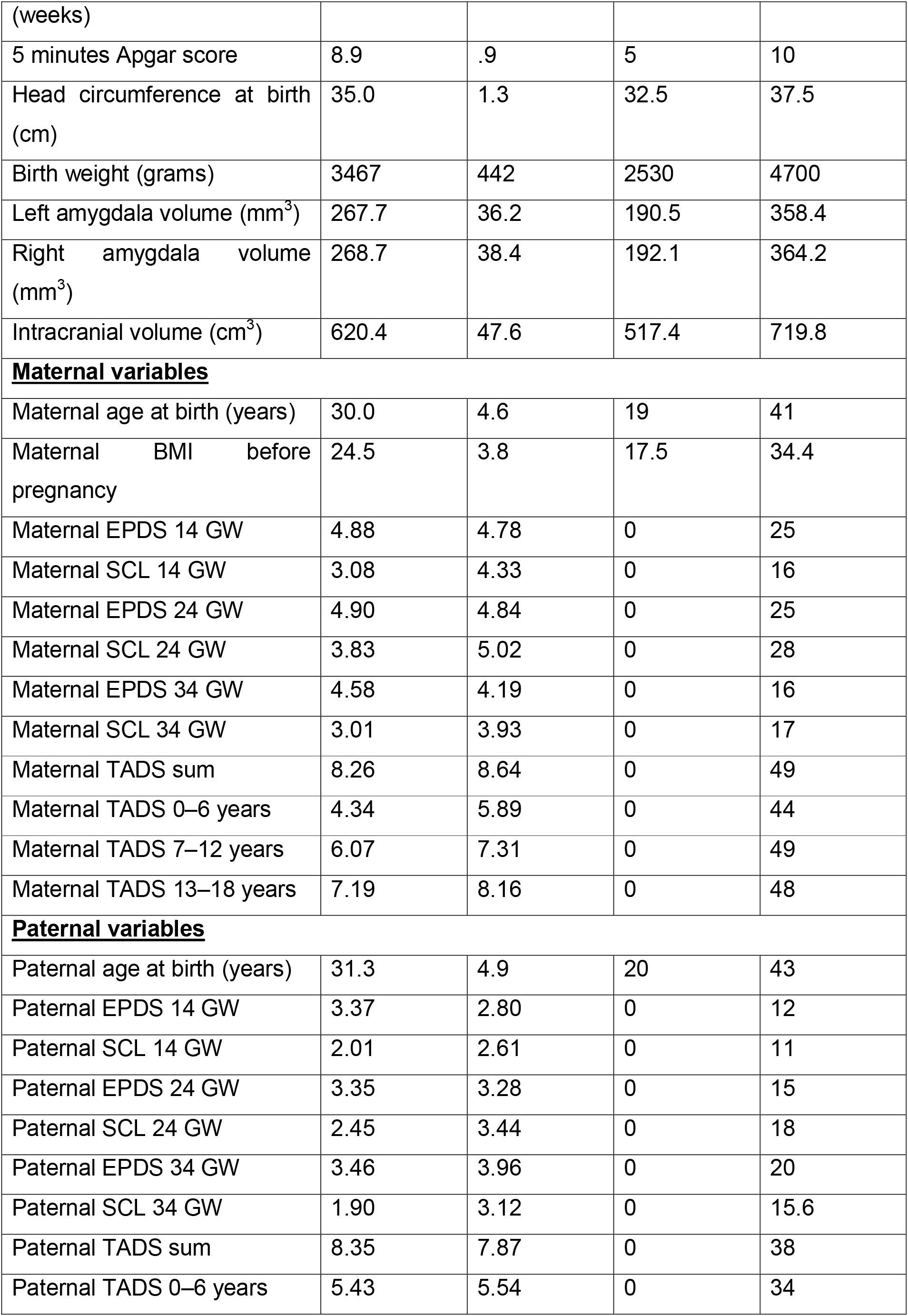
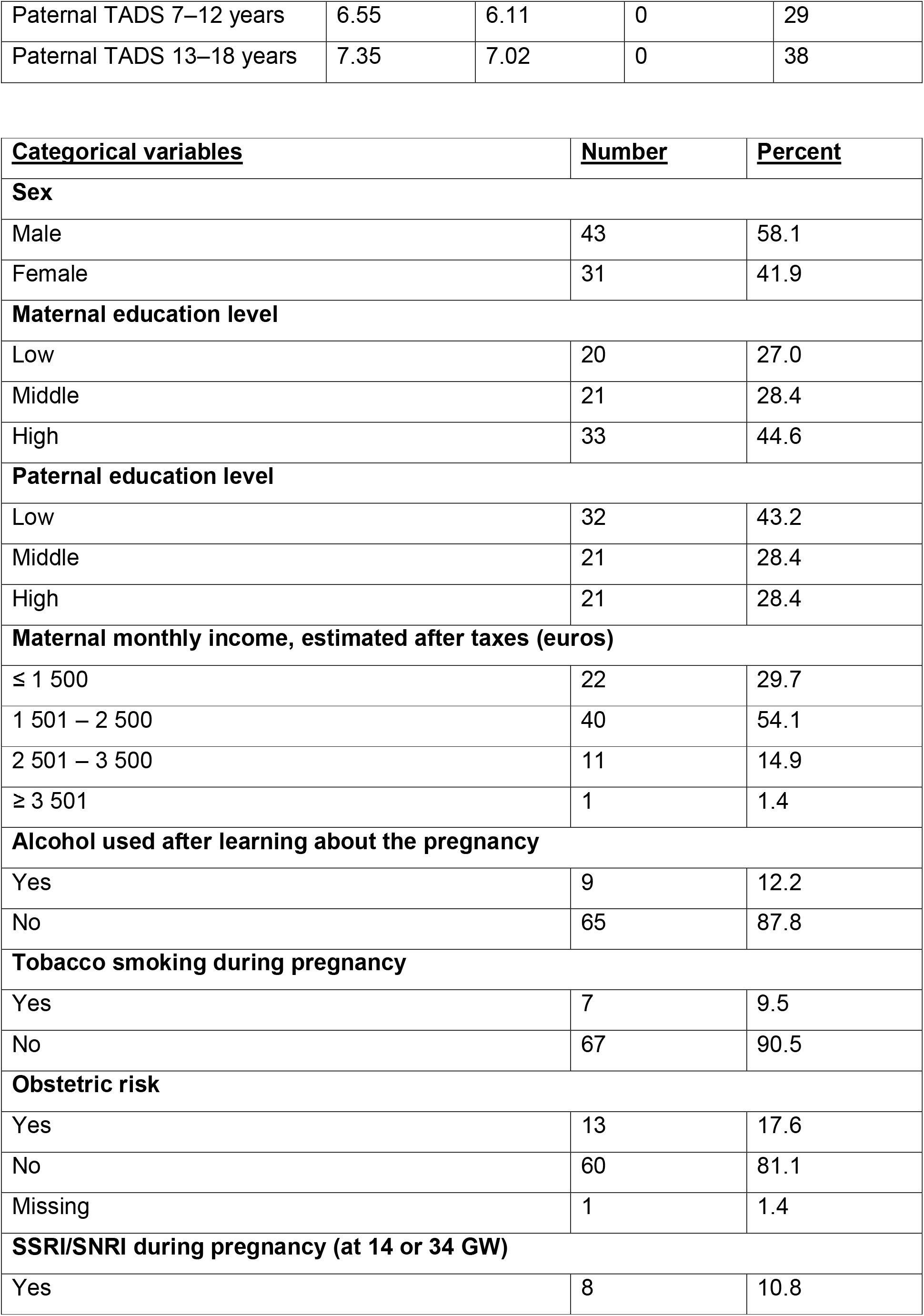
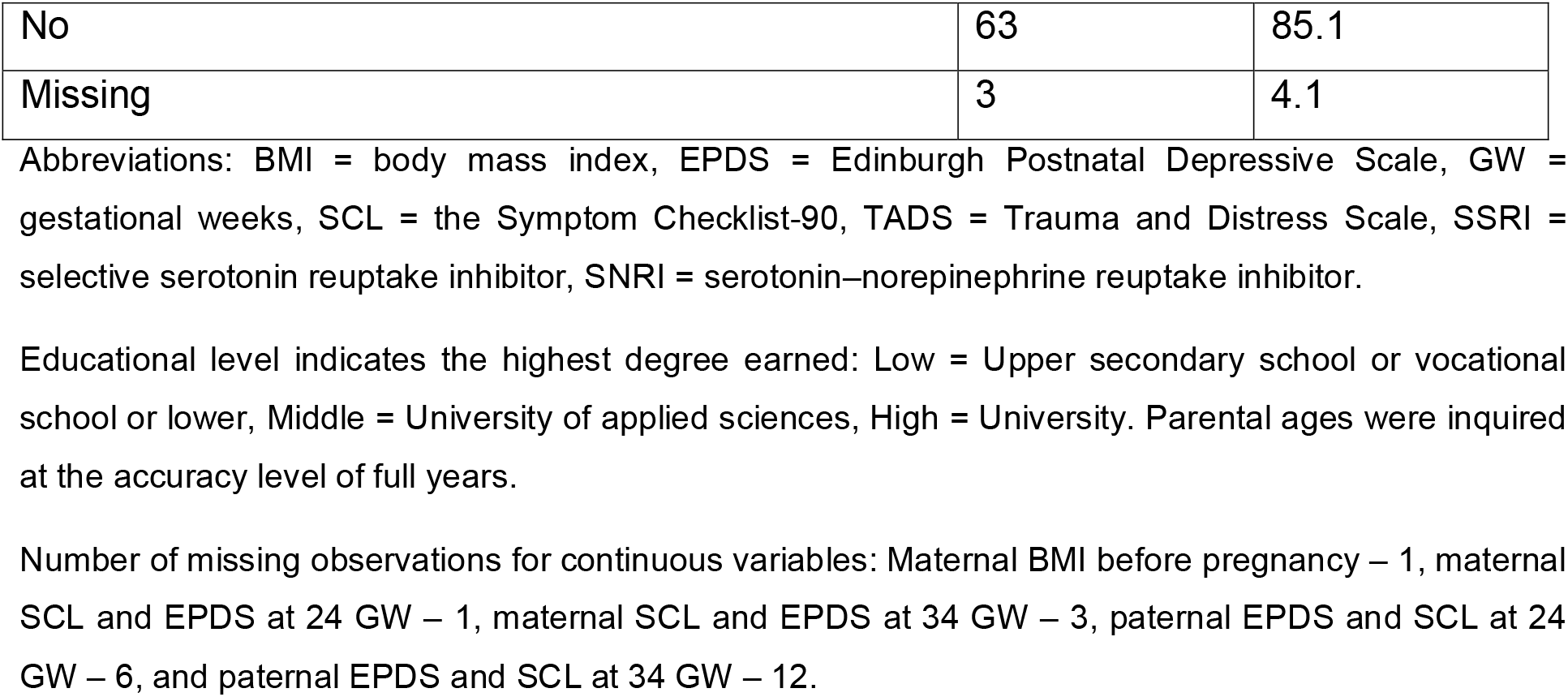
Participant demographics (N = 74).

#### Trauma and Distress Scale (TADS) questionnaire

Questionnaire data was collected at GW 14. TADS questionnaire includes five core domains: emotional neglect, emotional abuse, physical neglect, physical abuse, and sexual abuse, which were calculated according to Salokangas et al. (2016). The sum scores for these five core domains were calculated separately for traumatic events at 0–6, 7–12, and 13–18 years of age. The TADS sum score was calculated by taking the answer indicating most severe exposure from any timepoint for each question.

#### MRI acquisition

A detailed description of the scanning protocol is provided in our previous publication (Copeland et al. 2021; Lehtola et al. 2019). All the scans were performed at the same site with the same scanner and scanning parameters. Participants were scanned without anesthesia with a Siemens Magnetom Verio 3□T scanner (Siemens Medical Solutions, Erlangen, Germany). The 40-minute imaging protocol included an axial PD-T2-TSE (Dual-Echo Turbo Spin Echo) and a sagittal 3D-T1-MPRAGE (Magnetization Prepared Rapid Acquisition Gradient Echo) sequence, both with isotropic 1.0□mm^3^ voxels and whole brain coverage. Repetition time (TR) time of 12 070□ms and effective Echo time (TE) of 13□ms and 102□ms were used in PD-T2 TSE sequence to produce both PD-weighted and T2-weighted images from the same acquisition. Total number of 1cmm thick slices was 128. TR of 1900□ms, TE of 3.26□ms, and inversion time (TI) of 900cms were used for 3D-T1-MPRAGE sequence. The number of slices was 176.

All brain images were assessed for incidental findings by a pediatric neuroradiologist. If found, parents were given a follow-up opportunity with a pediatric neurologist. Developmental status was age appropriate by the age of two years for all participants. The incidental findings (intracranial hemorrhages, N□=□12, 6.7% of the participants with any imaging data) have been found to be common and clinically insignificant in previous studies (Rooks et al. 2008; Whitby et al. 2004). Intracranial hemorrhages were deemed not to affect volumetric estimates of interest and thus, did not exclude the individuals from the current study (Kumpulainen et al. 2020). All the brain images were checked visually by multiple researchers for motion artifacts.

#### Segmentations of amygdala

The amygdala segmentation has been described in detail in Acosta et al. (2020). A template was constructed based on the 125 good quality MRI scans (for the T1 data) using the methods described in (Fonov et al. 2011). The template was then warped to the center of 21 clusters of the data, and these 21 warped versions of the template were then manually labelled. Those 21 labelled warped templates were then used to create a consensus labelling was created using majority voting. This produced the final labelled template, which was then used to produce a template library for each subcortical structure. Segmentation into left and right amygdalae was done for each subject using a label-fusion-based labelling technique based on Coupé et al. (Coupé et al. 2011) and further developed by Weier et al. (2014) and Lewis et al. (2019). Finally, a mask of intracranial structures was automatically extracted for use in estimating intra cranial volume (ICV).

#### Code availability

Custom computer codes were used for data analysis. The core of the code to construct the library, and also to perform the fusion labeling can be found in https://github.com/NIST-MNI/nist_mni_pipelines/blob/master/iplScoopFusionSegmentation.py. Please contact the corresponding author for other code related questions.

#### Characterization of the final sample, risk of bias, and chosen confounders

The final sample for the current study, included all infant participants with neuroimaging data that had been successfully processed, and paternal and maternal data (including their TADS scores and relevant demographics); there were 74 such participants. There was missing information, with 1–12 values missing in the following parental variables (see Table 1 footnotes): maternal and paternal EPDS, pre-pregnancy BMI, use of SSRIs and SNRIs, and obstetric risk. For statistical analyses, missing information was imputed using mean (continuous variables) or mode (categorical variables) values of the variables. A flowchart of how the final sample was reached is shown in Figure 1.

**Figure 1.**
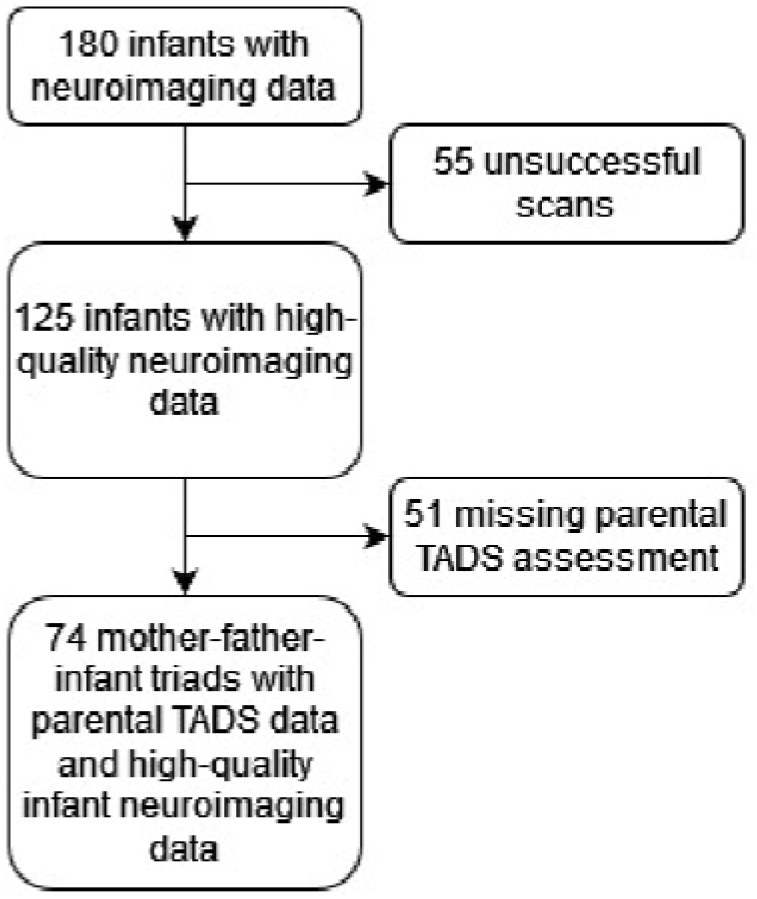
A flowchart describing the formation of the final sample of this study.

Risk of bias assessment is as follows: the exposed and non-exposed parents were drawn from the same population (low risk of bias), TADS questionnaire probes CME retrospectively in a single time point (intermediate risk of bias), the outcome of interest, infant amygdala volumes, were not present at the start of the study or at the time of exposure (low risk of bias), the statistical models controlled for important confounders (low risk of bias).

#### Potential maternal confounding factors

were also assessed (Pulli et al. 2019). Maternal alcohol use during pregnancy was inquired at GWs 14 and 34. At GW 14, 19 mothers reported using alcohol either weekly (N = 3), once or twice a month (N = 6), more rarely (N = 10), or did not specify the frequency (N = 1). The amount ranged from 0.5 to 7 units at a time (mean 2.3). Five reported continuing alcohol use to some extent after learning about the pregnancy. At GW 34, eight mothers reported using alcohol weekly (N = 1), once or twice a month (N = 1), or more rarely (N = 6). The amount ranged from 0.5 to 1 unit at a time. For the sake of statistical analyses, those who reported use at some point after learning about the pregnancy were considered the alcohol exposed group (N = 9). Data on tobacco exposure was combined from our three pregnancy questionnaires and from the National Institute for Health and Welfare (www.thl.fi) register. Seven mothers reported smoking tobacco at some point during the pregnancy. Eight mothers reported using SSRI/SNRI medication at either GW 14 or 34 (three participants had missing data in at least one timepoint). Additionally, obstetric risk was formalized in a binary value indicating the presence of one or more of the following risk factors extracted from the medical record: hypertension, diabetes, severe anaemia, severe infection, or vaginal bleeding. In our sample, 11 mothers had gestational diabetes mellitus and three had gestational hypertension (without significant proteinuria). In total, 13 mothers were considered to have obstetric risk (one had both diabetes and hypertension).

#### Statistical analyses

Regression models were calculated using R Statistical Software (v4.2.1; R Core Team 2022) in RStudio (version 2022.07.1+554). Zero order Spearman’s correlations between all covariates included in the regression analyses were made using JASP version 0.16.1.0 (JASP Team 2022) and are presented in Supplementary Table 2.

We first used linear regression models to test the associations between amygdala volume and parental TADS sum scores. The other parent’s TADS sum score, infant age from conception, infant age from birth, infant sex, infant intracranial volume (ICV; cm^3^), maternal education (three classes based on the highest degree earned: 1) Low = Upper secondary school or vocational school or lower; 2) Middle = University of applied sciences; 3) High = University), maternal EPDS score at GW 24, and maternal pre-pregnancy BMI were included as covariates in the analysis. Thereafter, we conducted a backward stepwise regression with Akaike information criterion (AIC) to optimize the variables left in the final regression model. Additionally, we performed sensitivity analyses where potential maternal confounding factors (alcohol exposure, tobacco exposure, SSRI/SNRI exposure, and obstetric risk) were added into the model one at a time.

Second, in analyses where we found a significant correlation between a parental TADS sum score and infant amygdala volume, we also performed a stepwise regression analysis (both forward and backward) with AIC using parental TADS scores from 0–6, 7–12, and 13–18 years separately (otherwise the same covariates as in the primary regression model).

For all three reported linear regression models, no multicollinearity between other covariates was detected (variance inflation factor (VIF) for all variables was between 1.07 and 1.25), and the correlation of residuals were > .99 for all models. The normal distribution of residuals was confirmed visually using Q-Q plots, scatter plots, and histograms. Additionally, Shapiro-Wilk test was calculated for all models and the p values ranged from .42 to .71, supporting the idea that the residuals were normally distributed.

We conducted a post hoc analysis to examine whether maternal EPDS at GW 24 acts as a mediator between maternal TADS sum score and infant left amygdala volume. We used the R package robmed (Alfons, Ateş, and Groenen 2022) in this analysis because EPDS and TADS scores were not normally distributed. The robmed package uses the robust MM-estimator and fast-and-robust bootstrap methodology. The model was first applied with all the same covariates as in the regression model, but the model did not convergent. Consequently, only the covariates that were significant for left amygdala volume in Supplementary Table 2 were included in the model. All reported estimates are according to the bootstrapped values (5000 resamples).

We did not perform formal corrections for multiple comparisons and report raw p values throughout the manuscript. The Bonferroni correct p value for statistical comparisons over left / right amygdala volumes is p < 0.025.

## Results

### Amygdala volume and parental TADS sum scores

For left amygdala volume, the significant predictors were ICV (β = .492, p < .001), age from birth (β = -.217, p = .043), and maternal TADS sum score (β = .223, p = .045). For right amygdala volumes, the significant predictors were ICV (β = .610, p < .001) and maternal education level (middle compared to other groups, β = .302, p = .003). For the full regression model, please see Table 3. In the sensitivity analyses there were no major differences to the main statistical model, although, in the sensitivity model that included maternal prenatal smoking the association between left amygdala volume and maternal TADS sum score was weaker (β = .191, p = .055).

### Amygdala volume and exploratory analyses with parental TADS scores at different ages

The exploratory analyses that included timing of the parental childhood maltreatment were conducted only for the left amygdala volume as there were no associations between TADS scores and right amygdala volume. The significant predictors were ICV (β = .474, p < .001), age from birth (β = -.243, p = .019), maternal TADS score from 13–18 years (β = .282, p = .008), and paternal TADS score from 0– 6 years of age (β = .247, p = .014). For the full regression model, please see Table 4.

### Is the effect of maternal TADS scores on infant left amygdala volume mediated by maternal EPDS score at GW 24?

Between maternal TADS sum score and left amygdala volume, there were marginally significant total (B = .778, p = .094) and direct (B = .844, p = .065) effects, but no significant mediation effect with maternal EPDS score at GW 24 (B = -.066, 95% confidence interval from -.463 to .051). The overall regression was not statistically significant (R^2^ = .305, F = 1.655, p = .157). In a model without covariates, the findings were similar, and there were no statistically significant associations (p > .05 for all effects). There were also no significant effects after adding paternal TADS sum score into the model.

## Discussion

We found that maternal CME associated positively with infant amygdala volume while the positive association for the paternal CME was only marginally significant (Table 2). In the exploratory analyses, we used age ranges for the timing of the CME (ages ranges: 0–6, 7– 12, and 13–18 years) and included all three time points from both parents in a stepwise regression model. We found that maternal CME between 13–18 years of age associated positively with infant left amygdala volumes. We also found that paternal CME between 0– 6 years of age associated positively with the infant left amygdala volumes. Similar associations were not observed for the right amygdala. These results add to the results of three previous studies linking maternal CME with amygdala volumes, although the studies have mixed findings (Demers et al. (2022), Moog et al. (2018), Khoury et al. (2021). To our knowledge, this is the first study to explore the associations between infant amygdala volume and paternal CME. The amygdala is one of the key brain structures associated with both early life exposures and later psychiatric health, which makes it crucially important to elucidate both the underlying mechanisms and the later relevance of these associations in future studies.

**Table 2.**
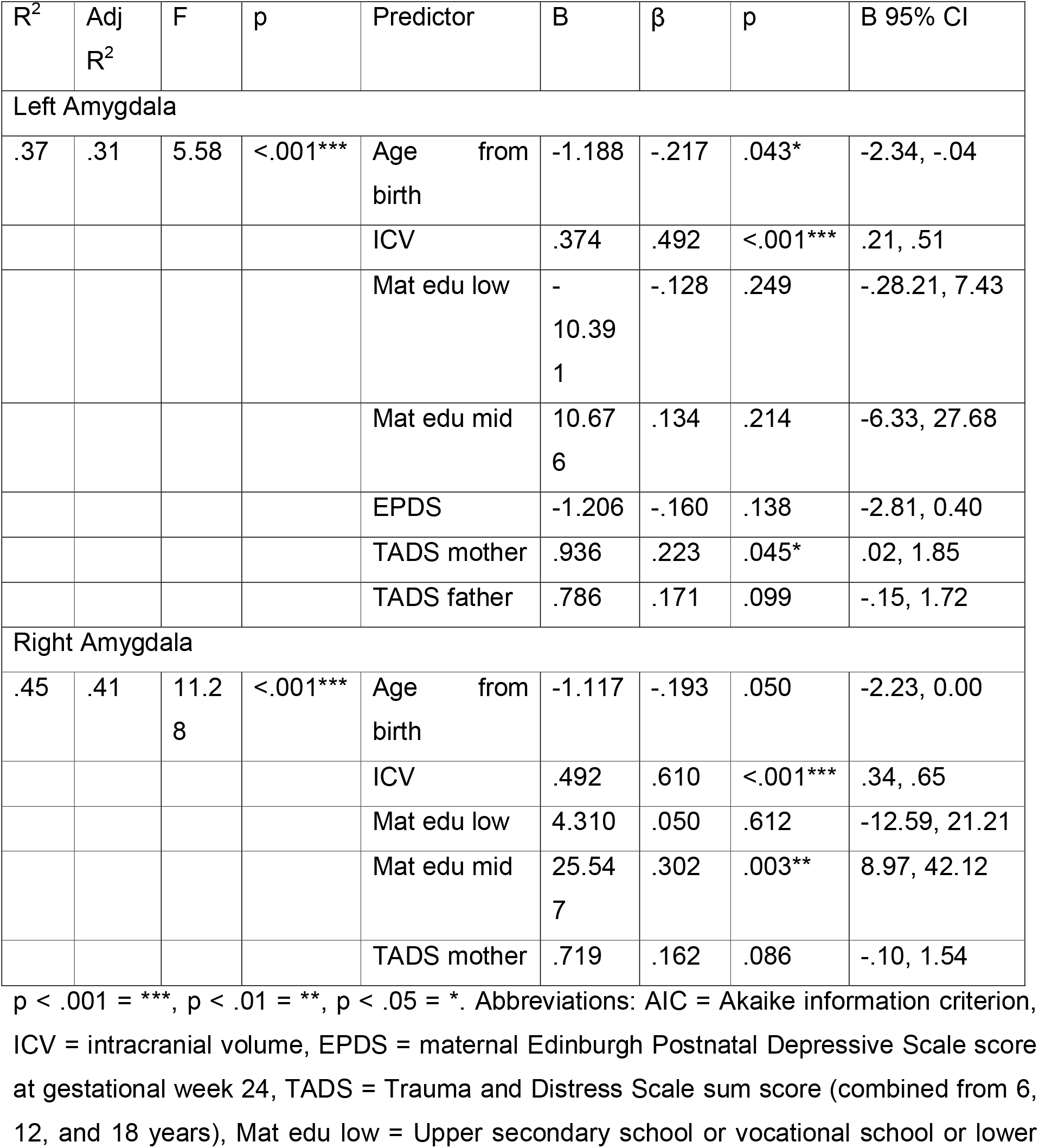
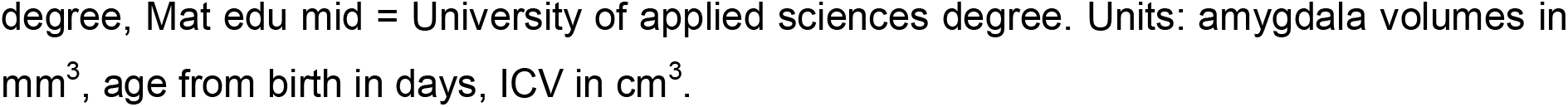
Stepwise backward linear regression model with AIC for infant amygdala volumes. TADS sum scores for mother and father.

**Table 3.** Stepwise forward and backward linear regression model with AIC for infant left

| R<sup>2</sup> | Adj<br>R<sup>2</sup> | F | p | Predictor | B | β | p | B 95% CI |
| --- | --- | --- | --- | --- | --- | --- | --- | --- |
| .43 | .37 | 7.05 | <.001*** | Age from birth | -1.327 | -.243 | .019* | -2.43, -.22 |
|  |  |  |  | ICV | .360 | .474 | <.001*** | .21, .51 |
|  |  |  |  | Mat edu low | -7.735 | -.095 | .365 | -24.68, 9.21 |
|  |  |  |  | Mat edu mid | 14.160 | .177 | .089 | -2.22, 30.54 |
|  |  |  |  | EPDS | -1.195 | -.159 | .125 | -2.73, .34 |
|  |  |  |  | TADS mother 13–18 years | 1.251 | .282 | .008** | .33, 2.17 |
|  |  |  |  | TADS father 0–6 years | 1.613 | .247 | .014* | .34, 2.89 |

amygdala volumes. Parental TADS scores from 0–6, 6–12, and 12–18 years separately.
p < .001 = \*\*\*, p < .01 = \*\*, p < .05 = \*. Abbreviations: AIC = Akaike information criterion, ICV = intracranial volume, EPDS = maternal Edinburgh Postnatal Depressive Scale score at gestational week 24, TADS = Trauma and Distress Scale score, Mat edu low = Upper secondary school or vocational school or lower degree, Mat edu mid = University of applied sciences degree. Units: amygdala volumes in mm<sup>3</sup>, age from birth in days, ICV in cm<sup>3</sup>.

### Potential intergenerational mechanisms of Maternal and Paternal maltreatment

During gestation CME-related physiological changes may be conveyed directly to the developing offspring via the feto–placental signaling. In a physiological sense, this is the only direct pathway of intergenerational transmission, and it continues throughout the pregnancy (Moog et al. 2022). Notably, in this study, depressive symptoms during gestation were measured and included into the analyses but they were not significantly associated with amygdala volumes. We lack the biological samples to elucidate concrete biological mechanisms, and these remain crucially important to map out in future studies. Possible mechanisms include changes in HPA axis / cortisol signaling and low-grade inflammation in the mother (Moog et al. 2018). In contrast to mothers, fathers do not have a direct connection with the fetus during gestation but can have an indirect influence through maternal wellbeing. None of the prior studies investigated paternal childhood trauma exposure in association with infant brain development. We have found that paternal CME has associations to infant gray matter volumes (Tuulari et al. 2022) and white matter integrity (Karlsson et al. 2020). In the current study, paternal effects were also positive but much statistically weaker than maternal associations. Paternal associations could be mediated via epigenetic changes in paternal germ line, but this remains to be verified in humans. Finally, direct genetic effects from parents to offspring are always possible so that parents with CME may be more likely from family conditions with higher psychiatric morbidity and infant amygdala volumes can reflect shared, genetic vulnerability.

### Contradictory findings among studies addressing parental early life adversity

Three previous studies have explored the associations between maternal CME and infant amygdala volume development, with mixed findings. Two of these studies found a negative association between maternal ACEs (including CME and household dysfunction): Demers et al. (2022) with bilateral amygdala and Khoury et al. (2021) in a sample of 4–24-month-old infants they found a negative association between maternal CME and infant amygdala volume, and the effect was stronger in infants at the older end of the age range, which implies that the associations in early infancy can lead to altered developmental trajectories or that the postnatal environment is contributing to the effect. Moog et al. (2018) did not find any associations. We found a positive association between maternal CME and left amygdala volume, which does not replicate any of the prior findings and is contradictory to the results obtained by Demers et al. (2022). As such the variable findings are very common in studies of intergenerational exposures. For instance, similarly mixed findings exist for “maternal prenatal distress exposures” (Supplementary Table 1). The sample sizes of all extant studies, including ours, have been small (< 100), have used variable measures of ACEs / CME, MRI methods, and may have limited comparability because of different societal contexts. Parental reports of early adversity are obtainable retrospectively in any ongoing study and if such measures will be obtained there are good opportunities to replicate and extend these initial findings.

### Comparison of measures of parental early adversity

In addition to TADS questionnaire (Salokangas et al. 2016) that was used in the current study, there are other well-validated CME questionnaires, such as the childhood trauma questionnaire (CTQ; Bernstein et al. 1994, 1997, 2003). The core domains of TADS and CTQ (emotional and physical neglect and abuse, sexual abuse) are very similar. Adverse Childhood Experiences (ACEs) Questionnaire (Felitti et al. 1998), a 10-item survey used to measure childhood trauma. The questionnaire assesses ten types of childhood trauma. Five are personal: physical abuse, verbal abuse, sexual abuse, physical neglect, and emotional neglect. Five are related to family members: parent with alcohol-related problems, a mother/caretaker who is a victim of domestic violence, a family member in jail, a family member diagnosed with psychiatric disorder, and the disappearance of a parent through divorce, death, or abandonment. More detailed assessment of early life adversity can be obtained with interviews such as the Childhood Trauma Interview (CTI; Thabrew, De Sylva, and Romans 2012).

### Limitations

Our study has some limitations. First, data on childhood maltreatment was collected in adulthood, potentially resulting in recall error. However, this measurement has been shown to be valid, reliable, and clinically relevant (Luutonen et al. 2013; Salokangas et al. 2016). Recall error also precludes closer analyses that probe longitudinal CME, questionnaires have limited capability in differentiating CME-related acute vs. chronic stress, and it is not possible to identify precise causes of stress reactions that have occurred. Second, it is difficult to pinpoint the exact role of CME in individual development as well as the mechanism responsible for the intergenerational transmission. In this study, depressive symptoms during pregnancy did not mediate this connection. Third, the sample is very ethnically homogenous, meaning the results are not necessarily generalizable to other ethnicities. Similarly, the sample hasn’t been selected/enriched based on the severity of the symptoms (e.g., posttraumatic stress disorder patients), meaning that the findings are not generalizable to clinical populations. Finally, the sample size is relatively small although comparable to prior studies.

## Conclusions

In this study, we found a positive association between infant left amygdala volume and maternal CME while controlling for covariates. The effect of paternal CME on left amygdala volume was not statistically significant. The maternal association was not mediated by maternal depressive symptoms during pregnancy. In an analysis exploring the timing of the CME, left amygdala volumes were positively associated with maternal CME at 12–18 years and paternal CME at 0–6 years of age. There are few studies on the intergenerational effects of CME, and more are needed. In future studies, it is especially important to include information of parental CME. Furthermore, identification of sensitive time periods for both mothers and fathers is an important goal for future studies.

## Acknowledgements

We would like to warmly thank all FinnBrain families that participated to the study. We would also like to thank the research team: Satu Lehtola for her help in data collection, Maria Lavonius for her help in recruiting the participants, Jani Saunavaara for implementing the MRI sequences, Riitta Parkkola for reviewing the MR images for incidental findings.

## Author contributions

JJT and EPP planned the analytical approach and performed the data analyses, lead writing of the manuscript. ELK contributed significantly to writing and editing the initial draft of the manuscript. JDL contributed to the planning of the MRI sequences, developed the methods, and carried out the brain segmentation. LP contributed to statistical analyses. LK and HK planned and funded the MRI measurements, established the FinnBrain Birth Cohort and built the infrastructure for carrying out the study. All authors participated in conceptualizing the study, writing the manuscript and accepted the final version.

## Funding

Jetro J. Tuulari

Sigrid Jusélius Foundation; Emil Aaltonen Foundation; Finnish Medical Foundation; Alfred Kordelin Foundation; Juho Vainio Foundation; Turku University Foundation; Hospital District of Southwest Finland; State Grants for Clinical Research (ERVA); Orion Research Foundation, Signe and Ane Gyllenberg Foundation.

Elmo P. Pulli

Päivikki and Sakari Sohlberg Foundation; Juho Vainio Foundation.

Eeva-Leena Kataja

Academy of Finland. Grant number: #346790; Signe and Ane Gyllenberg Foundation; Turku University Foundation

Linnea Karlsson

Academy of Finland, Grant/Award Number: #325292; State Grants for Clinical Research (ERVA) (P3654); Brain and Behavior Research Foundation, YI Grant #1956; Signe and Ane Gyllenberg Foundation

Hasse Karlsson

Academy of Finland; Turku University Foundation; Hospital District of Southwest Finland; State Grants for Clinical Research (ERVA);

