## Supplemental Table 1 for "Parental childhood maltreatment associates with offspring left amygdala volume at early infancy"

Supplementary Table 1. A literature review of previous studies exploring the associations between infant/child amygdala volume and parental ACEs and on a closely related intergenerational exposure maternal prenatal distress.

| Study | Subjects | Distress assessment | Confounders | Controlling pre-/postnatal distress | Other areas inspected | Amygdala results | Other results |
|---|---|---|---|---|---|---|---|
| **Exposure: Maternal ACEs** | | | | | | | |
| (Demers et al., 2022) | N = 85<br>(44 females)<br>42–51 weeks postconceptional age<br><br>The Care Project | The ACE Questionnaire | ○ Sex<br>○ Birth weight percentile<br>○ Parity<br>○ Obstetric complications during the index pregnancy<br>○ Income-to-needs ratio<br>○ Intracranial volume<br>○ Postconceptional age at scan | No | Hippocampus | ↓ Higher maternal ACEs associated with lower bilateral amygdala volumes | No significant association between maternal ACEs and bilateral hippocampal volume |
| (Khoury et al., 2021) | N = 57<br>4–24 months<br><br>the Harvard MIND (Mother Infant Neurobiological Development) study | The 10-item ACE questionnaire and<br>the 75-item MACE scale | ○ Sex<br>○ Gestational age at birth (in weeks)<br>○ Family income<br>○ Maternal education | EPDS at 4 months postpartum | TBV<br>GMV<br>WMV<br>Cerebrospinal fluid volume<br>Hippocampus | ↓ Infant age moderated the association between maternal ACEs and right amygdala volume, such that ACEs were associated with lower volume at older ages. | Maternal ACEs associated with lower infant total brain volume and gray matter volume, with no moderation by infant age |

| | | | | | | | |
|---|---|---|---|---|---|---|---|
| (Moog et al., 2018) | N = 80 newborns,<br><br>Development, Health and Disease Research Program | The Childhood Trauma Questionnaire | ○ SES<br>○ Obstetric complications<br>○ Obesity<br>○ Recent interpersonal violence<br>○ Gestational age at birth<br>○ Sex<br>○ Postnatal age at MRI | Pre- and early postpartum stress | Intracranial volume<br>Gray matter volume<br>White matter volume<br>Cerebrospinal fluid<br>Hippocampus | No associations with amygdala | Maternal CM exposure was associated with lower child intracranial volume, which was primarily due to a global difference in cortical gray matter |
| **Exposure: Prenatal distress** | | | | | | | |
| (Groenewold et al., 2022) | N = 124<br>(63 females)<br>3 weeks<br><br>The Drakenstein Child Health Study (DCHS) | BDI-II (GW 28-32) | ○ Sex<br>○ Age at scan<br>○ Intracranial volume (ICV)<br>○ Clinic site<br>○ Maternal age<br>○ Education level<br>○ Alcohol and tobacco exposure | No | Hippocampus<br>Thalamus<br>Caudate<br>Putamen<br>Pallidum | ↑ Right Amygdala | ↑ Bilateral Hippocampus<br>↑ Bilateral Caudate<br><br>Interaction by child sex (females with stress exposure had larger hippocampi) |
| (Lautarescu et al., 2021) | N = 221<br>(107 females)<br>Term-equivalent age, M = 42.20 GWs | (Prenatal) STAI and Stressful life events | ○ GA<br>○ PMA<br>○ SES<br>○ Sex<br>○ Days on total parenteral nutrition | | Frontal lobe<br>Temporal lobe<br>Hippocampus<br>Thalamus | No associations between prenatal stress and infant brain. | No associations between prenatal stress and infant brain. |

| | | | | | | | |
|---|---|---|---|---|---|---|---|
| (Moog et al., 2021) | N = 86<br>5–64 postnatal days, mean: 25.8 ± 12.9 | (Prenatal) PSS | ○ Infant sex<br>○ Maternal SES<br>○ Maternal smoking during pregnancy<br>○ Maternal obstetric complications during pregnancy | Sensitivity analyses: use of psychotropic medication and alcohol use as covariates | Hippocampus | Prenatal stress was not associated with amygdala volumes | ↓ Left hippocampal volume |
| (Wu et al., 2021) | N = 101<br>Neonates<br>(38–47 gestational weeks) | PSS, EPDS, STAI<br>(23-40 GWs) | | Prenatal medication | Cortical grey matter and white matter<br>Deep grey matter<br>Cerebellum<br>Brainstem<br>Hippocampus | ↓ Bilateral amygdala volumes | No associations with prenatal stress. |
| (Acosta et al., 2020) | N = 28 (14 females)<br>4 years | EPDS 14, 24, 34 GWs | ○ Maternal prenatal medication<br>○ Alcohol and/or tobacco nicotine exposure<br>○ Education<br>○ BMI<br>○ Gestational complications<br>○ Previous miscarriages and/or abortions<br>○ Maternal childhood trauma | Pre- and postnatal general anxiety (SCL-90)<br>Postnatal depressive symptoms (EPDS) | - | ↓ Higher 2nd trimester EPDS significantly related to smaller right amygdalar volumes in the whole sample.<br><br>↓ Higher 3rd trimester EPDS associated with significantly smaller right amygdalar volumes in boys compared to girls | - |

| | | | | | | | |
|---|---|---|---|---|---|---|---|
| | | | ○ Child's birth weight<br>○ Gestational age | | | | |
| (Lehtola et al., 2020) | 123 newborns (69 males, 54 females)<br>2–5 weeks | SCL-90<br>EPDS;<br>A combination variable | ○ Age at scan<br>○ Sex<br>○ Maternal prenatal medication (CNS-affecting medication)<br>○ Substance use (alcohol or tobacco)<br>○ Maternal education<br>○ Gestational diabetes<br>○ Birth complications (asphyxia, CRP, respirator treatment, 5 min Apgar score under five) | - | Hippocampus | ↓ Relative volumes of both the left and right amygdala and the interaction term of sex by SCL-90 + EPDS (from GW 24) showing a more negative association with amygdalar volumes in males than in females. | No significant associations were found between SCL + EPDS sum scores and hippocampal volumes in the sex-interaction model. |

| | | | | | | | |
|---|---|---|---|---|---|---|---|
| (Lee et al., 2019) | N = 167 neonates<br>N = 199 (age 4.5 y) | (Prenatal) EPDS<br>GW 26;<br>EPDS median (> 9) split, high vs low | ○ Age at MRI<br>○ Maternal ethnicity<br>○ Education | BDI-II and EPDS at 3 months, 1 year, 2 years, 3 years, and 4.5 years postpartum | Amygdala–cortical circuits | In the “high prenatal depression group”, a positive correlation between amygdala volume and insula thickness in girls.<br><br>At 4.5 years, negative association between amygdala volume and inferior frontal thickness. | - |
| (Wang et al., 2018) | N = 161 neonates<br><br>GUSTO | EPDS<br>26 GWs | ○ Age at MRI<br>○ Total brain volume<br>○ Maternal education<br>○ Gender<br>○ Maternal ethnicity | | Hippocampus<br>Cortical grey matter and white matter<br>CSF | ↓ A trend-significant association between prenatal stress and amygdala volumes moderated by the FKBP5 gene | ↓ Association with (lower) right hippocampal volume (moderated by genetic variants in FKBP5).<br><br>A trend-significant association between prenatal stress and cortical thickness moderated by FKBP5. |
| **Exposure: Pre- and postnatal distress** | | | | | | | |
| (Hidalgo et al., 2022) | N = 2993<br>10 years<br><br>Generation R | (Prenatal) The Social Readjustment | ○ Child sex<br>○ Age at MRI<br>○ Total intracranial volume | | Total brain volume<br>Cortical grey and cerebral | No relationship between maternal prenatal adversity and brain volumes. | Prenatal adversities not related to any brain outcome. |

| | | | | | | | |
|---|---|---|---|---|---|---|---|
| | | Rating Scale (SRRS)<br>20-25 GWs -> "prenatal adversities score"<br><br>(Postnatal) Childhood adversities (24 events occurring during the 0-10 years interviewed by the mother) | ○ Maternal ethnicity<br>○ Highest household education (for SES)<br>○ Prenatal alcohol and tobacco exposure | | white matter volumes<br>Cerebellar volume<br>Hippocampus | Postnatal adversities were not related to the amygdala or hippocampal volumes. | Postnatal adversities were not related to hippocampal volumes.<br><br>Per each additional childhood adverse event, the total brain volume was 0.07 standard deviations smaller (SE = 0.02, $p = 0.001$), with differences in both grey and white matter volumes. |
| (Donnici et al., 2021) | N = 54<br>(25 females)<br>3–7 years<br><br>Alberta Pregnancy Outcomes and Nutrition (APrON) study | SCL-90R<br>(Prenatal)<br>2nd trimester (17 GWs);<br>(Postnatal)<br>12 weeks postpartum | ○ Child's gestational age at birth<br>○ Birth weight<br>○ Household income<br>○ Age<br>○ Sex | Pre-/postnatal maternal depression (EPDS) | - | No associations for prenatal anxiety.<br><br>↑ Postnatal anxiety associated with increased amygdala volume (<u>but association disappeared when controlling for total brain volume</u>) | - |
| (Wen et al., 2017) | N = 342<br>4.5 years<br><br>GUSTO | (Prenatal)<br>EPDS<br>26 GW; | ○ Age at MRI<br>○ Maternal ethnicity<br>○ Maternal education | | - | ↑ Prenatal maternal depressive symptoms were associated with | - |

| | | | | | | | |
|---|---|---|---|---|---|---|---|
| | | Postnatal EPDS at 3 months, BDI-II at 1, 2, 3 and 4.5 years | ○ Total brain volume<br>○ Sex | | | larger right amygdala volume in girls, but not in boys.<br><br>↑ Postnatal maternal depressive symptoms were associated with higher right amygdala FA in the overall sample and girls, but not in boys | |
| (El Marroun et al., 2016) | N = 654 (327 females)<br><br>6–10 years<br><br>The Generation R | Brief Symptom Inventory (BSI) | ○ Sex<br>○ Age<br>○ Maternal age<br>○ Education<br>○ Smoking during pregnancy<br>○ Birth weight<br>○ Gestational age<br>○ Child emotional and behavioral problems | Maternal depressive symptoms at 3 years | Total brain volume Hippocampus | Prenatal maternal (or paternal) depressive symptoms were not associated with differences in amygdala.<br><br>Depressive symptoms at 3 years were not associated with amygdala. | Prenatal maternal (or paternal) depressive symptoms were not associated with differences in total brain volume or bilateral hippocampus.<br><br>Depressive symptoms at 3 years were not associated with total brain volume or bilateral hippocampus. |

Abbreviations: ACE = Adverse Childhood Experiences, MACE = Maltreatment and Abuse Chronology of Exposure, EPDS = Edinburgh Postnatal Depression Scale, TBV = total brain volume, GMV = gray matter volume, WMV = white matter volume, SES = socioeconomic status, MRI = magnetic resonance imaging, CM = childhood maltreatment, BDI = The Beck Depression Inventory, GW = gestational week, STAI = The State-Trait Anxiety Inventory, GA = gestational age, PMA = postmenstrual age, PSS = Perceived Stress Scale, BMI = body mass index, SCL-90 = The Symptom Checklist 90, CNS = central nervous system, CRP = C-reactive protein, GUSTO = Growing Up in Singapore Towards healthy Outcomes, CSF = cerebrospinal fluid, FA = fractional anisotropy.
