## Supplemental Table 2 for "Parental childhood maltreatment associates with offspring left amygdala volume at early infancy"

**Correlation Table**

| Variable | | Left Amygdala | Right Amygdala | Left Amygdala ICV corrected | Right Amygdala ICV corrected | ICV | Paternal TADS sum score | Maternal TADS sum score | Age from conception | Age from birth | Sex | Maternal pre-pregnancy BMI | Maternal EPDS at gestational week 24 | Maternal education, high vs. other | Maternal education, low vs. other |
|---|---|---|---|---|---|---|---|---|---|---|---|---|---|---|---|
| 1. Left Amygdala | Spearman's rho | — | | | | | | | | | | | | | |
| | p-value | — | | | | | | | | | | | | | |
| 2. Right Amygdala | Spearman's rho | 0.652 | — | | | | | | | | | | | | |
| | p-value | < .001 | — | | | | | | | | | | | | |
| 3. Left Amygdala ICV corrected | Spearman's rho | 0.818 | 0.397 | — | | | | | | | | | | | |
| | p-value | < .001 | < .001 | — | | | | | | | | | | | |
| 4. Right Amygdala ICV corrected | Spearman's rho | 0.506 | 0.859 | 0.534 | — | | | | | | | | | | |
| | p-value | < .001 | < .001 | < .001 | — | | | | | | | | | | |
| 5. ICV | Spearman's rho | 0.474 | 0.549 | -0.074 | 0.073 | — | | | | | | | | | |
| | p-value | < .001 | < .001 | 0.529 | 0.534 | — | | | | | | | | | |
| 6. Paternal TADS sum score | Spearman's rho | 0.231 | 0.158 | 0.150 | 0.100 | 0.191 | — | | | | | | | | |
| | p-value | 0.048 | 0.179 | 0.201 | 0.394 | 0.103 | — | | | | | | | | |
| 7. Maternal TADS sum score | Spearman's rho | 0.066 | 0.072 | 0.072 | 0.060 | 0.030 | 0.212 | — | | | | | | | |
| | p-value | 0.578 | 0.542 | 0.541 | 0.613 | 0.798 | 0.070 | — | | | | | | | |
| 8. Age from conception | Spearman's rho | 0.123 | 0.248 | -0.152 | -0.005 | 0.500 | 0.079 | 0.170 | — | | | | | | |
| | p-value | 0.295 | 0.033 | 0.197 | 0.968 | < .001 | 0.505 | 0.147 | — | | | | | | |
| 9. Age from birth | Spearman's rho | -0.016 | 0.035 | -0.233 | -0.188 | 0.331 | 0.080 | 0.071 | 0.449 | — | | | | | |
| | p-value | 0.893 | 0.766 | 0.045 | 0.109 | 0.004 | 0.499 | 0.546 | < .001 | — | | | | | |
| 10. Sex | Spearman's rho | -0.220 | -0.287 | -0.010 | -0.122 | -0.344 | -0.037 | -0.174 | -0.027 | -0.169 | — | | | | |
| | p-value | 0.060 | 0.013 | 0.935 | 0.299 | 0.003 | 0.757 | 0.139 | 0.820 | 0.150 | — | | | | |
| 11. Maternal pre-pregnancy BMI | Spearman's rho | 0.019 | -1.185e-4 | -0.073 | -0.065 | 0.117 | 0.004 | 0.073 | 0.041 | 0.109 | 0.037 | — | | | |
| | p-value | 0.870 | 0.999 | 0.535 | 0.584 | 0.322 | 0.972 | 0.537 | 0.731 | 0.355 | 0.757 | — | | | |
| 12. Maternal EPDS at gestational week 24 | Spearman's rho | -0.039 | 0.091 | -0.051 | 0.069 | 0.048 | 0.114 | 0.385 | 0.246 | 0.121 | -0.076 | 0.028 | — | | |
| | p-value | 0.743 | 0.443 | 0.666 | 0.557 | 0.686 | 0.332 | < .001 | 0.035 | 0.306 | 0.519 | 0.812 | — | | |
| 13. Maternal education, high vs. other | Spearman's rho | -0.041 | -0.197 | -0.045 | -0.268 | 0.024 | 0.011 | 0.060 | -0.017 | 0.048 | 0.010 | 0.028 | 0.055 | — | |
| | p-value | 0.726 | 0.093 | 0.702 | 0.021 | 0.842 | 0.923 | 0.612 | 0.889 | 0.686 | 0.935 | 0.813 | 0.641 | — | |
| 14. Maternal education, low vs. other | Spearman's rho | 0.177 | 0.115 | 0.101 | -0.009 | 0.171 | -0.043 | -0.021 | 0.076 | 0.141 | -0.038 | 0.060 | 0.122 | 0.546 | — |
| | p-value | 0.132 | 0.328 | 0.391 | 0.942 | 0.145 | 0.717 | 0.856 | 0.519 | 0.232 | 0.746 | 0.613 | 0.301 | < .001 | — |

Abbreviations: ICV = intracranial volume, TADS = Trauma And Distress Scale, BMI = body mass index, EPDS = Edinburgh Postnatal Depressive Scale
